# Rapid Antibody Fragment Production and Binding Analysis Using Cell-Free Protein Synthesis Combined with Fluorescence Correlation Spectroscopy

**DOI:** 10.1101/2025.08.05.668779

**Authors:** Shakiba Nikfarjam, Chao Liu, Emma J. Laurence, Adam N. Laurence, Jennifer L. Chlebek, Dante P. Ricci, Steven A. Hoang-Phou, Ted A. Laurence, Matthew A. Coleman

## Abstract

This study investigates the efficient development and production of single-chain variable fragments (scFvs) and antibody fragments (Fabs) using an *E. coli*-based cell-free protein synthesis system. Validation of the methodology was performed using a fluorescence correlation spectroscopy (FCS)-based assay to determine binding equilibrium constants (*K*_D_) between antibodies and the receptor binding domain (RBD) of SARS-CoV-2 Spike protein. An initial assessment employed two conventionally cell-produced anti-RBD antibodies. To optimize cell-free production, folding strategies were developed to enhance the solubility and yields of scFvs, including a two-stage refolding protocol that successfully recovered active proteins from misfolded precipitates. Fab fragments were also produced and characterized, with their binding properties analyzed to assess functionality. This study highlights the potential of cell-free systems for the rapid and efficient production of functional antibody fragments. The integration of advanced techniques, such as FCS-based kinetic measurements, underscores the versatility and applicability of cell-free platforms for antibody development and high-throughput screening. These findings offer a promising avenue for accelerating therapeutic antibody research and production.

**SIGNIFICANCE of WORK:** This study highlights the transformative potential of integrating cell-free protein synthesis (CFPS) with fluorescence correlation spectroscopy (FCS) for the rapid and scalable production and characterization of functional antibody fragments. By leveraging an *E*. *coli*-based CFPS platform, we successfully designed and optimized the production of single-chain variable fragments (scFvs) compared to Fab fragments, addressing some common challenges of low solubility and yield. The development of a cost-efficient two-stage refolding strategy further enhanced the scalability and functionality of scFvs, enabling higher recovery of active proteins from the unfolded state. Additionally, FCS provided a sensitive, rapid method for accurately quantifying antigen-antibody binding kinetics across a range of affinities. This work establishes CFPS and FCS as versatile and complementary tools, offering a robust framework for accelerating antibody fragment development, particularly in time-sensitive scenarios like infectious disease outbreaks.

## INTRODUCTION

The COVID-19 pandemic has highlighted the urgent need for rapid and efficient methods of protein production, particularly for the development of potential vaccines and therapeutic antibodies. Traditional protein expression systems, such as transient transfection in mammalian cells, often require significant time and resources, which can delay the discovery and development of critical therapeutics (1–3). To address this challenge, cell-free protein synthesis (CFPS) has emerged as a promising approach for the rapid and flexible production and screening of proteins. CFPS systems, such as those based on Escherichia coli (*E. coli*) and Chinese hamster ovary (CHO) lysates, eliminate the need for cell growth and maintenance, enabling faster screening coupled with multiple production cycles compared to conventional cell-based methods (4–8). CFPS retains the protein-producing functionality of the cell in a lysate format, allowing for *in vitro* protein synthesis without the constraints of cell viability (5,9,10). While *E. coli*-based lysates are widely used for their cost-effectiveness and high yields (9), eukaryotic lysates, such as those derived from CHO cells, are advantageous for producing complex proteins requiring post-translational modifications (9–12). However, eukaryotic systems can be less efficient and more expensive than their prokaryotic counterparts (9).

Antibodies play a pivotal role in therapeutic and diagnostic applications, with single-chain variable fragments (scFvs) and fragments of antibodies (Fabs) offering significant advantages due to their smaller size and ease of engineering and stability (11–14). While intact antibodies, such as IgG, are commonly used in research and therapeutics (11), smaller fragments are often preferred for high-throughput screening and production due to their simplified structure (15). There are multiple methods for design and production of antibody fragments. These include variable heavy domain of heavy chain (VHH), which is binding portion of heavy chain-only antibodies (15), scFv, which is the smallest recombinant antibody fragment that still contains the antigen-binding domain of the original antibody (16), and Fab, which is the ‘antigen binding fragment’ that includes the entire antigen-binding domain of an antibody (17), but are structured more similarly to IgG than scFv(18). Despite their advantages, traditional cell-based expression systems often suffer from low yields and solubility issues, which can limit their scalability and utility for rapid screening (19–22). Cell-free protein synthesis (CFPS) has emerged as a powerful alternative (23), enabling efficient and scalable production of functional antibody fragments (23–25). Furthermore, optimization strategies, such as chaperone supplementation, redox tuning, and post-expression refolding protocols, have been shown to enhance solubility and facilitate the recovery of properly folded and functional proteins from aggregated or misfolded intermediates (26–28).

In addition to rapid protein production, binding kinetics are critical for therapeutic and diagnostic development. Conventional techniques for determining binding rates, such as surface plasmon resonance (SPR) (29–31), enzyme-linked immunosorbent assays (ELISAs) (25,26), or more recent methods like Bio-Layer Interferometry (BLI) (31,32) can be time-consuming and often require substantial amounts of purified protein. To overcome these limitations, Fluorescence Correlation Spectroscopy (FCS) offers a powerful alternative that enables real-time, high-throughput binding measurements with minimal sample requirements, even in complex mixtures such as crude lysates. (33–38). FCS detects fluorescently labeled molecules as they diffuse through a highly confined detection volume of around 1 femtoliter. Upon binding, molecular complexes exhibit slower diffusion due to their increased hydrodynamic radius, allowing for precise determination of binding equilibrium constants (*K*_D_), which inherently reflect the underlying association and dissociation rates (38, 39). Beyond binding measurements, FCS offers several advantages that make it uniquely powerful: it enables quantification of molecular concentrations down to the single-molecule level, provides insights into molecular weight and oligomerization states based on diffusion dynamics, and supports real-time kinetic studies in complex mixtures, including crude cell-free reactions. Moreover, FCS permits multiplexed or site-specific analyses with minimal material consumption, making it exceptionally well-suited for evaluating the expression, folding, and functionality of antibody fragments produced via CFPS (38,41–44).

This study integrates CFPS and FCS to highlight a versatile platform for the rapid production and quantitative screening of several different affinity reagents. This is demonstrated using an *E. coli*-based CFPS system for production of human-relevant scFvs and Fab fragments, providing protocols for overcoming low solubility as well as low yields. A two-stage refolding protocol was developed to recover insoluble scFv, while Fab fragments were directly expressed and characterized. The binding properties of these fragments were analyzed using FCS, which provided precise and high-throughput assessment of antigen-antibody interactions Whereas conventional SPR workflows typically require multiple days for protein preparation and kinetic measurements—often spanning 2–7 days overall (45,46)—our CFPS-to-FCS pipeline consolidates the complete workflow into a single day. This substantial time reduction makes the pipeline particularly advantageous for rapid screening applications. These findings highlight the potential of CFPS and FCS as complementary tools for accelerating therapeutic antibody development, particularly in time-sensitive scenarios such as infectious disease outbreaks.

## MATERIAL and METHODS

### Cell-free expression systems and optimization

We used CFPS protocols as previously described by Liu et al. (47). Briefly, ClearColi BL21(DE3) cells were grown in 2x YT media with 1% NaCl, induced with 1 mM IPTG, harvested, and lysed via sonication. Lysates were clarified through centrifugation, underwent a 30-minute run-off reaction at 37 °C, and were stored at −80 °C. For CFPS, reactions (1 mL scale) were prepared by combining lysate (25% final volume), plasmid DNA, and a reaction mix containing 1.2 mM ATP, 0.86 mM GTP/UTP/CTP, 34 µg/mL folinic acid, 170 µg/mL E. coli tRNA, 2 mM of each amino acid (except glutamic acid), 0.33 mM NAD, 0.27 mM acetyl-CoA, 1.5 mM spermidine, 1 mM putrescine, 175 mM potassium glutamate, 10 mM ammonium glutamate, 2.7 mM potassium oxalate, 10 mM magnesium glutamate, and 33 mM phosphoenolpyruvate (PEP). In figure 1, the workflow of E. coli lysate preparation is illustrated.

**Figure 1.**
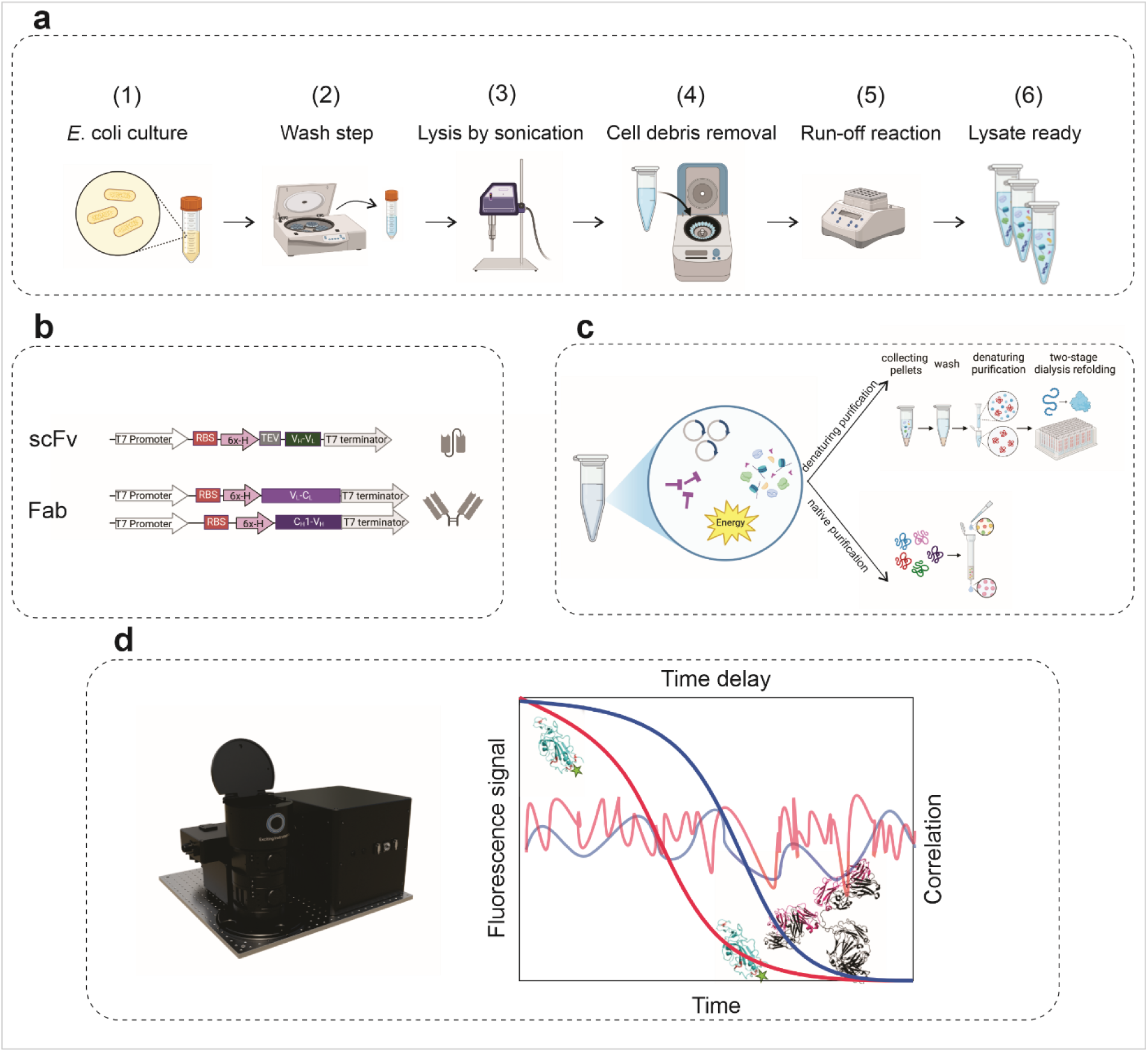
Workflow for lysate preparation to the production of purified antibody fragments. This figure provides an overview of the process used for cell-free antibody fragment production and purification and screening. **(a)** Lysate preparation: The **E. coli** lysate system was prepared as the foundation for the cell-free protein synthesis (CFPS) platform following the general scheme shown. **(b)** Construct design: DNA templates for the production of antibody fragments were designed and cloned into the Pivex2.3d plasmid, enabling efficient transcription and translation. This illustration highlights the modularity of the DNA constructs used for generating both single-chain variable fragments (scFvs) and Fab fragments combined with prokaryotic translational regulatory sequences. **(c)** Protein synthesis and purification: Antibody fragments were synthesized using the CFPS system, followed by downstream purification steps to isolate and prepare the functional proteins for characterization and further analysis. The top arrow shows denaturing purification of scFvs, and the lower arrow shows the native state purification of Fab fractions. (d) Commercial confocal FCS instrument (EI-FLEX), time trace of fluorescence signal. Red signal and corresponding correlation curve show signal from labeled RBD before binding to the antibody, faster fluctuations of laser intensity is due to smaller size and fast diffusion of unbound RBD. Blue signal and the corresponding correlation curve, shows the slower movement of the labeled RBD upon binding to the antibody.

### Construct design

The protein-coding DNA sequence for antibody fragments 2130-parental and 80R-2.59 were synthesized, and the remaining variants were generated using PCR mutagenesis. The gene was subsequently cloned into the PiveX2.3d vector. Details of the scFv and Fab construct sequences are provided in the supplementary materials. For the Fab fragments, two constructs were tested. Both constructs shared the same Fab heavy chain, but one of the two Fab constructs included a FLAG tag in addition to the His-tag on the light chain.

### Protein expression and purification

#### IgG Production

Full-length 2130 antibody was produced as an IgG in transiently transfected CHO cells as described in Desautels et al.(48) The S309 IgG was purchased from ATUM.

#### scFv Production and Purification

For scFvs, expression and solubility were monitored in cell-free reactions without the addition of disulfide enzymes or chaperones. Most scFv variants exhibited low solubility in the *E. coli* cell-free lysate, with some being completely insoluble (Supplementary Material). Solubility significantly improved upon the addition of 4 μM DsbC and 5 μM DnaK (PUREfrex, Supplementary Material). To further enhance the purification yields of scFvs for binding assays, a modified two-stage refolding protocol was applied to refold scFvs from insoluble cell-free precipitates. The use of CFPS markedly reduced the overall time required for production, purification, and refolding of scFvs, cutting the process from four days to two. Additionally, the protein precipitates from CFPS were less contaminated with cell debris and impurities, eliminating the need for extensive multi-step detergent washes prior to resuspension in guanidine hydrochloride. The cell-free reaction consisted of 25% lysate, 25% reaction buffer, 15-20 μg/mL plasmid, appropriate additives, and ultrapure water. Reactions were incubated overnight at 30 °C and 800 RPM in a thermomixer. After incubation, the reactions were centrifuged at 10,000 × g to remove insoluble material and subjected to Ni-NTA purification according to the supplier’s protocol (Roche 08778850001). For denaturing purification, 1 mL of reactions were incubated overnight in a 24-well plate on a thermomixer, increasing surface area to enhance the production of insoluble proteins. The reactions were then centrifuged at 18,000 × g to collect protein precipitates, which were washed twice with Tris-buffered saline (pH 8). The washed pellets were resuspended in denaturing buffer (6 M guanidine-HCl, 100 mM Tris (pH 8), 10 mM imidazole) and loaded onto a Ni-NTA column under denaturing conditions (20 mM sodium phosphate (pH 6.5), 0.5 M NaCl, 0.2 M imidazole, and 8 M urea). The protein was eluted in two fractions: the first with 250 mM imidazole and the second with 500 mM imidazole. The eluted denatured scFvs were refolded using a two-step dialysis process. In the first step, proteins were dialyzed overnight in 2 L of 100 mM Tris (pH 8.6), 2 mM glycine, 1 mM EDTA, 1 mM L-cysteine, and 2.5 mM urea. The second step involved dialysis for 3 hours in PBS, followed by an additional exchange of PBS for 2 hours. The final yield of purified scFvs was approximately 200 uL of 0.5 mg/mL for each variant.

#### Fab Production and Purification

The use of CFPS for the expression and purification of Fab fragments was straightforward. The 1 mL reactions produced high yields of soluble and active Fab fragments without requiring disulfide-forming additives or chaperones. Fab cell-free reactions were centrifuged at 8,000 × g for 10 minutes and subsequently subjected to Ni-NTA purification. The final yields of purified soluble Fab fragments were approximately0.5 mL of 0.3-0.4 mg/mL.

#### Expression, Purification, and Fluorescent Labeling of RBD

The receptor-binding domain (RBD) of the SARS-CoV-2 spike protein was expressed using the Expi293 transient transfection system according to the manufacturer’s protocol. The expression construct was cloned into the pSF-CMV vector, which includes a CMV promoter for high-level expression in mammalian cells. The encoded RBD sequence contains an N-terminal leader peptide to facilitate secretion and a C-terminal Avi-His tag for affinity purification. The expected molecular weight of the expressed protein, including all tags, is 34 kDa. Following expression, the supernatant was collected and purified using Ni-NTA affinity chromatography, followed by size-exclusion chromatography (SEC) for further refinement. The final purified RBD had a concentration of 1.2 mg/mL. SDS-PAGE analysis confirmed purity, as shown in Figure 2a. For fluorescent labeling, the purified RBD was conjugated to Alexa Fluor 488 and Alexa Fluor 647 maleimide dyes via a site-specific labeling strategy targeting an a free cysteine in the upstream sequence before the His tag. To ensure selective reduction of the cysteine, a 50 µL aliquot of 35 µM RBD was incubated with a 2-fold molar excess of TCEP for 30 minutes at room temperature. Subsequently, a 2-fold molar excess of thiol-reactive Alexa (488 or 647) dye was added, and the reaction was incubated for 30 minutes at room temperature in the dark. The unreacted dye was removed by performing three consecutive rounds passes through using Zeba spin desalting columns (Thermo Fisher, Cat# 89882). The efficiency of fluorescent labeling was determined using dye absorbance measurements, with an estimated labeling efficiency of 30-40%. This optimized expression and labeling approach enabled the production of highly purified and fluorescently labeled RBD for downstream biophysical analyses.

**Figure 2.**
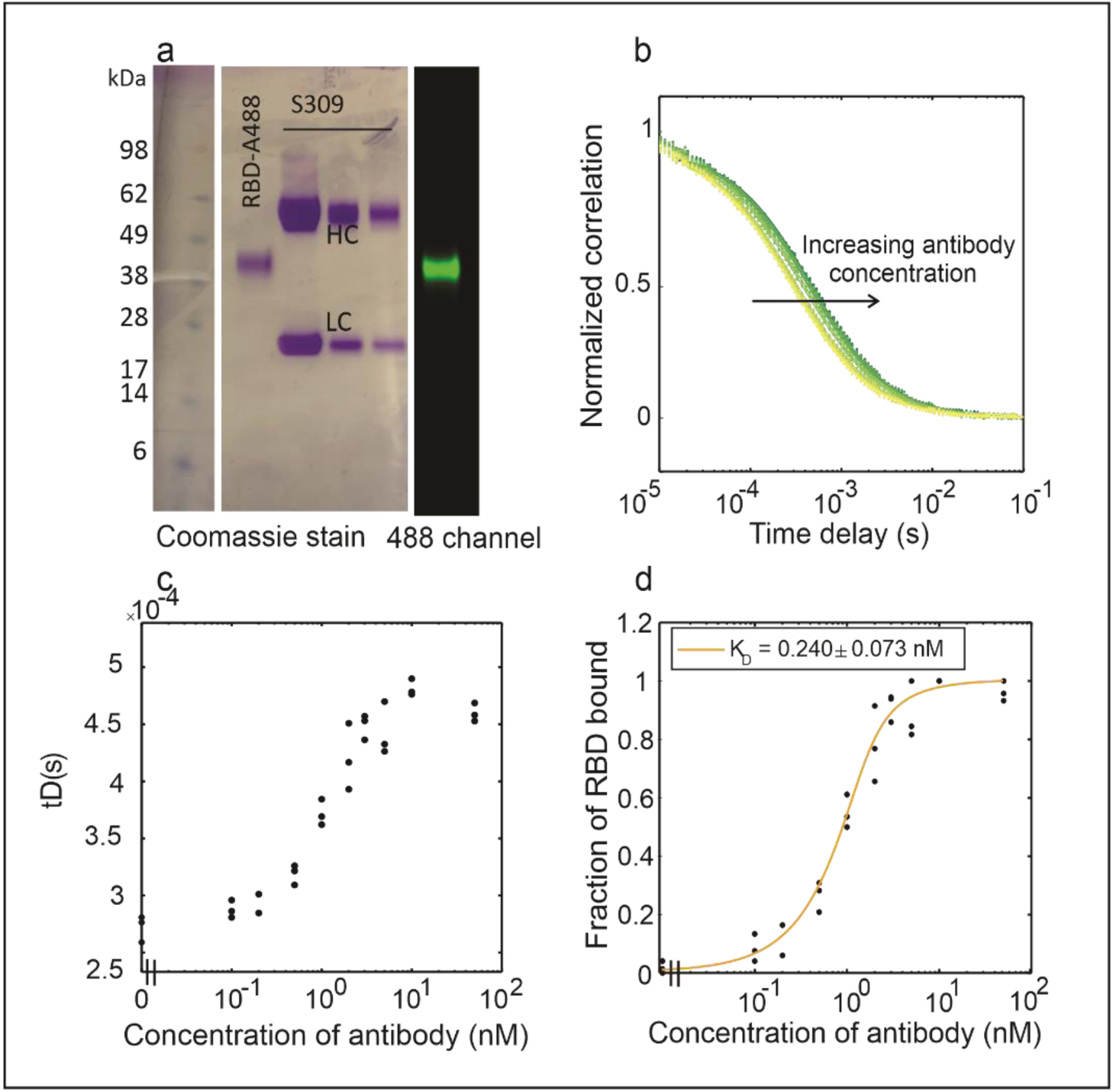
FCS measures sub-nanomolar binding affinity (*R*_*D*_) between S309 IgG and receptor-binding domain (RBD) of the SARS-CoV-2 spike protein. (**a**) SDS-PAGE gel of purified RBD (BQ.1-488) and three dilutions of S309 IgG. Lane 1: A488-labeled RBD, Lane 2: S309 IgG, lane 3: labeled RBD imaged at 488 nm wavelength confirming the proper labeling. HC: heavy chain; LC: light chain. **(b)** FCS correlation curves, data demonstrates a shift rightward with increasing IgG concentrations, indicating longer diffusion times due to RBD-antibody complex formation. **(c)** Diffusion time (Y-axis) increases as antibody concentrations rise (X-axis), plateauing when reaching saturation. **(d)** Fraction of bound RBD as a function of antibody concentration, fitted using a two-species binding model.

#### Fluorescence correlation spectroscopy (FCS) for Binding Measurements

Fluorescence Correlation Spectroscopy (FCS) is a powerful technique that analyzes temporal fluorescence intensity fluctuations of individual fluorophores as they diffuse through a femtoliter-sized confocal detection volume. This approach enables the determination of molecular diffusion coefficients, concentrations, and interactions. Similar to light scattering techniques, larger particles exhibit slower fluorescence fluctuations due to their extended residence time within the focal volume (49,50). The high sensitivity of FCS, capable of detecting as few as 1–10 molecules per confocal volume, allows for precise characterization of molecular binding dynamics (51,52). In an FCS experiment, fluorescently labeled molecules are excited by a focused laser, and emitted photons are collected by highly sensitive detectors. The temporal fluctuations in fluorescence intensity are analyzed using an autocorrelation function, which provides information on molecular concentration and diffusion dynamics. The autocorrelation function is defined as:

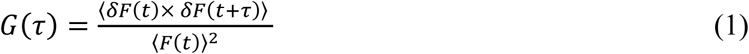

where *G*(τ) represents the second-order fluorescence intensity correlation as a function of the lag time τ, and 〈*F*(*t*)〉 is the average fluorescence intensity and δ*F* is the deviation (fluctuation) of the instantaneous signal from its mean.

The correlation data are fitted to extract the diffusion time τ_*D*_, which represents the characteristic time required for a molecule to traverse the confocal detection volume. For a single-component diffusion model, the correlation function is expressed as:

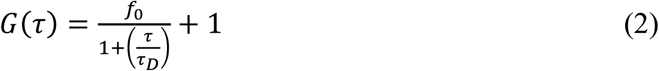

where *f*_0_ is the amplitude of the correlation function, and τ_*D*_ is the diffusion time.

To quantify antibody-antigen binding affinities, the autocorrelation data are fitted to a two-component diffusion model

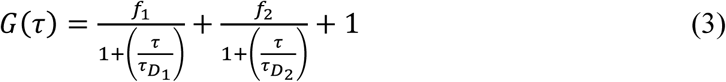

In this model, τ_*D*1_ represents the diffusion time of free receptor-binding domain (RBD), while τ_*D*2_ corresponds to the diffusion time of the antibody-bound RBD complex. By keeping τ_*D*1_ and τ_*D*2_ constant during fitting, the relative amplitudes *f*_1_ and *f*_2_ are extracted, corresponding to the amplitudes of the free and bound molecules, respectively.

The fraction of bound RBD is then determined using the equation (47):

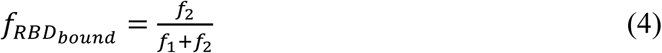

This method allows for the precise quantification of binding interactions by monitoring shifts in diffusion time as complexes form. By employing FCS, we can differentiate between free and bound species with high sensitivity, enabling real-time measurements of antibody-antigen interactions in solution. When the concentration of antigen, which in this study is fluorescently labeled RBD, is significantly lower than the dissociation constant (*K*_*D*_), the binding data can be approximated by a simple hyperbolic binding equation. Under these conditions, the fraction of bound RBD follows

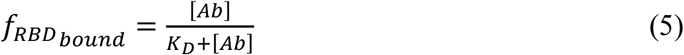

where [*Ab*] represents the antibody concentration. This equation allows for direct extraction of *K*_*D*_from the binding curve by plotting f_RBDbound_ as a function of [*Ab*] and fitting the data to the hyperbolic function. This approach is particularly useful in FCS measurements where the trace component (labeled RBD) is kept in low concentration, minimizing the effects of ligand depletion and simplifying the binding model.

To fit the binding curves obtained from full-length antibodies, we used the quadratic binding equation, which accounts for the non-negligible concentration of binding partners and is particularly important when ligand and receptor concentrations are comparable. The fluorescence correlation data were analyzed using the following form of the quadratic equation:

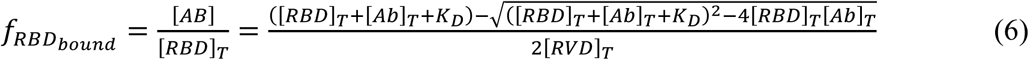

where *[RBD]_T_* is the total concentration of RBD, *[Ab]_T_* is the total concentration of the antibody (or antibody fragment), and *K_D_* is the dissociation constant. When *[RBD]_T_* was held constant and *[Ab]_T_* titrated, the equation was fitted to the measured binding curve (*f_RBD-bound_ vs [Ab]_T_*) with two fitting parameters: *[RBD]_T_* and *K_D_*. Conversely, when *[Ab]_T_* was held constant and *[RBD]_T_* titrated, the equation was fitted to *f_RBD-bound_ vs [RBD]_T_* with fitting parameters *[Ab]_T_* and *K_D_*. This approach enables robust determination of binding affinities without assuming a large excess of one binding partner.

By employing FCS, we can differentiate between free and bound species with high sensitivity, enabling real-time quantification of antibody-antigen interactions in the solution.

#### FCS instrumentation

The FCS measurements were performed using both a homebuilt system (47) and a commercially available instrument, the **EI-FLEX**. The detail of homebuilt FCS instrument used in this work is brought in detail in the previous work published buy Liu et, al (47). Briefly, the system was equipped with a multi-wavelength laser source (Stradus VersaLase 4, Vortran Laser) with excitation at 488 nm. The laser was fiber-coupled and directed into the detection volume via a 60× water immersion objective (UPlanApo/IR, 1.20 NA, Olympus), generating a confocal volume of approximately 1.6 fL. Fluorescence emission was collected through a 50 μm pinhole and split into red and green channels using an emission dichroic mirror (FF562-Di03-25×36, Semrock). Bandpass filters (FF01-609/54-25 and FF02-525/40-25, Semrock) further refined the emission before detection by two avalanche photodiodes (PD-050-CTD, Micro-Photon-Devices). The detection volume was calibrated using solutions of Atto 488 dye at known concentrations. The volume was determined by measuring fluorescence correlation curves for dye solutions at concentrations ranging from 500 pM to 100 nM. A linear fit to the inverse correlation amplitude versus concentration yielded an estimated detection volume of 1.5–1.8 fL, accounting for minor variations due to alignment and environmental factors. The FCS data collected for the full-length antibodies used the homebuilt system, and the data for the Fab fragments and scFvs was collected using EI-FLEX system, a commercially available FCS instrument. The instrument is equipped with 520 and 638 continuous wave lasers and two APD detectors.

## RESULTS

### FCS Measures Antibody-Antigen Binding Affinities

To validate FCS as a quantitative tool for measuring antibody-antigen interactions, we first tested its ability to detect binding using well-characterized monoclonal antibodies targeting the receptor-binding domain (RBD) of the SARS-CoV-2 spike protein. For this purpose, we selected S309 IgG, a broadly neutralizing antibody known to bind multiple RBD variants with nanomolar affinity(53). Figure 2a shows an SDS-PAGE gel confirming the high purity (>95%) of both the Alexa Fluor 488 (A488)-labeled RBD and the mammalian-expressed S309 IgG used in this validation experiment. For FCS-based binding measurements, we maintained a constant concentration (∼1 nM) of A488-labeled RBD while titrating increasing concentrations of S309 IgG (0–50 nM). The FCS correlation curves demonstrated a progressive rightward shift with increasing antibody concentrations (Figure 2b), corresponding to an increase in diffusion time as antibody-RBD complexes formed. This shift is expected, as larger molecular complexes diffuse more slowly through the confocal volume.

To quantify binding dynamics, diffusion times were extracted from the correlation curves by fitting the data to Equation 2 (see methods). Diffusion time measurements plateaued at high antibody concentrations (10 nM), consistent with binding saturation (Figure 2c). To determine binding affinity, we first fitted a two-species diffusion model to the correlation curves and obtained the amplitudes of unbound (*f*_₁_) and bound (*f*_₂_) RBD species using Equation 3. The fraction of the bound RBD was then calculated from Equation 4. By fitting the quadratic binding equation (Equation X) to the plot of fraction of bound RBDs vs antibody concentration, we determined that the S309 IgG and the WT-RBD of SARS-CoV-2 spike protein have a sub-nanomolar (*K*_*D*_ = 0.240 in this experiment) binding affinity (Figure 2d).

Following this validation, we applied FCS to compare the binding affinities of two monoclonal IgGs—wild-type2130 and S309—against three RBD variants: wild-type RBD, BQ.1, and BQ.1.1 (Figure 3). These RBD variants are different in several key mutations. The wild-type RBD sequence is the original SARS-CoV-2 spike receptor-binding domain, which binds ACE2 and is the target of many neutralizing antibodies. BQ.1 has four mutations and BQ.1.1 has five mutations (three of which are the same in both) relative to the wild-type RBD (54). The binding experiments were performed by titrating antibody concentrations against a constant concentration of fluorescently labeled RBD (1-3 nM). As antibody concentrations increased, the labeled RBD exhibited longer diffusion times, resulting in a rightward shift in the correlation curve as shown in Figure 2b, in which the correlation curves shift from yellow to green. The fraction of RBD bound is determined from two-species fits to the correlations (Equations 3 and 4) and plotted against antibody concentrations (Figure 3c). These binding curves were fitted to either a hyperbolic or quadratic function, enabling determination of fitted *K*_D_ values as indicated in table 1.

**Figure 3.**
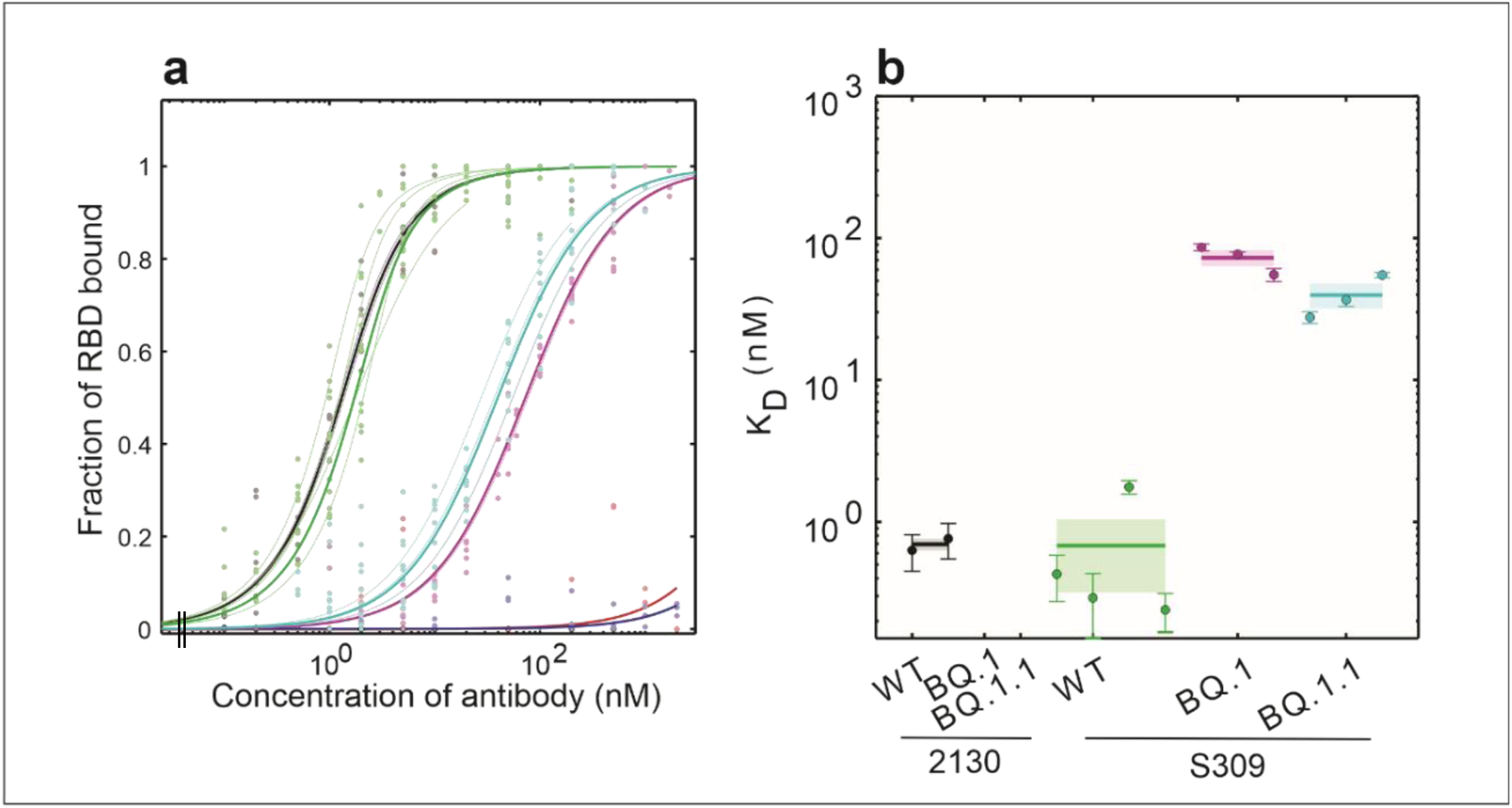
Differential Binding Affinities of Two IgGs Reveal Selective Recognition of RBD Variants. **(a)** A two species fit (eq. 3) to the correlation curves gives us the fraction of RBD-A488 bound to IgG at each IgG concentration. Each thin line represents one experiment (as described in Figure 2), and thick lines represent the means. See (b) for color legend. The blue and red curves are between 2130 IgG and WT RBD or BQ.1 RBD, respectively. **(b)** The dissociation binding constants extracted from the binding curves in (a). Each data point represents one thin curve (one experiment) from (a). Horizontal lines and shaded areas represent mean± s.e.m. Values are given in Table 1.

**Table 1.** Dissociation constants between antibodies and RBD variants of SARS-CoV-2 spike protein determined by FCS. Binding affinities between BQ.1, BQ1.1 and antibody 2130 were undetectable (no binding). Values are mean +-sem of multiple experiments.

| Antibody/RBD | WT | BQ.1 | BQ.1.1 |
| --- | --- | --- | --- |
| 2130 | 0.70±0.07 nM | - | - |
| S309 | 0.68±0.36 nM | 73±9 nM | 40±8 nM |

Wild-type 2130 IgG exhibited strong binding to wild-type RBD, with a sub-nanomolar *K*_*D*_of 0.70±0.07 nM. However, no detectable binding was observed between 2130 IgG and the BQ.1 and BQ.1.1 RBD variants suggesting that mutations in these variants resulted in loss of critical binding interactions. In contrast, S309 IgG displayed strong sub-nanomolar binding to wild-type RBD (0.68±0.36 nM) (table 1). However, S309 exhibited significantly reduced binding affinity to the BQ.1 and BQ.1.1 variants, with *K*_D_ values of 73±9 nM and 40±8 nM, respectively, suggesting that these RBD mutations impact the antibody’s binding epitope. These results underscore the versatility of S309 IgG in binding to RBD variants with key mutations. These findings also highlight the utility of FCS for characterizing binding interactions across a range of affinities and its potential for studying emerging viral variants.

The KD values that we determined by FCS are in reasonable agreement with values reported in the literature using conventional methods like surface plasmon resonance and biolayer interferometry (48,53,55–57). For example, previous affinity data showed the binding affinity between WT RBD and S309 IgG was ∼0.7 nM, which matches our FCS results (58). Together, these results validate FCS as a sensitive method for quantifying antigen-antibody binding affinities.

### Expression Screening of scFvs in *E. coli* Cell-Free System Reveals Solubility Constraints

To assess the efficacy of CFPS for producing single-chain variable fragments (scFvs), we utilized the commercial biotechrabbit RTS 100 *E. coli* HY Kit (Catalog #BR1400101). A total of six 2130 scFv derivatives (parental, 182, 183, 185, 186, 188) and five 80R scFv variants (003, 127, 44, 43, 32) were tested for expression in *E. coli*-based CFPS. To directly monitor scFv expression without the need for purification, we incorporated FluoroTect™ GreenLys *in vitro* Translation Labeling System (Catalog #L5001). This system enables real-time fluorescent visualization of newly synthesized proteins on SDS-PAGE gels by incorporating a BODIPY-labeled lysine-tRNA complex into nascent polypeptides (59).

Analysis of the CFPS reactions revealed low solubility across all expressed scFvs using the *E. coli* cell-free lysate. Specifically, the 2130 derivatives exhibited significantly lower solubility (<10% of the total protein expressed determined by densitometry analysis) than 80R variants (30-40% solubility) (Figure 4). This suggests the 80R scFv derivatives are about twice more soluble than the parental. The differences in solubility may be explained by intrinsic sequence-dependent folding properties which have previously been shown to influence solubility outcomes in CFPS (60–62).

**Figure 4.**
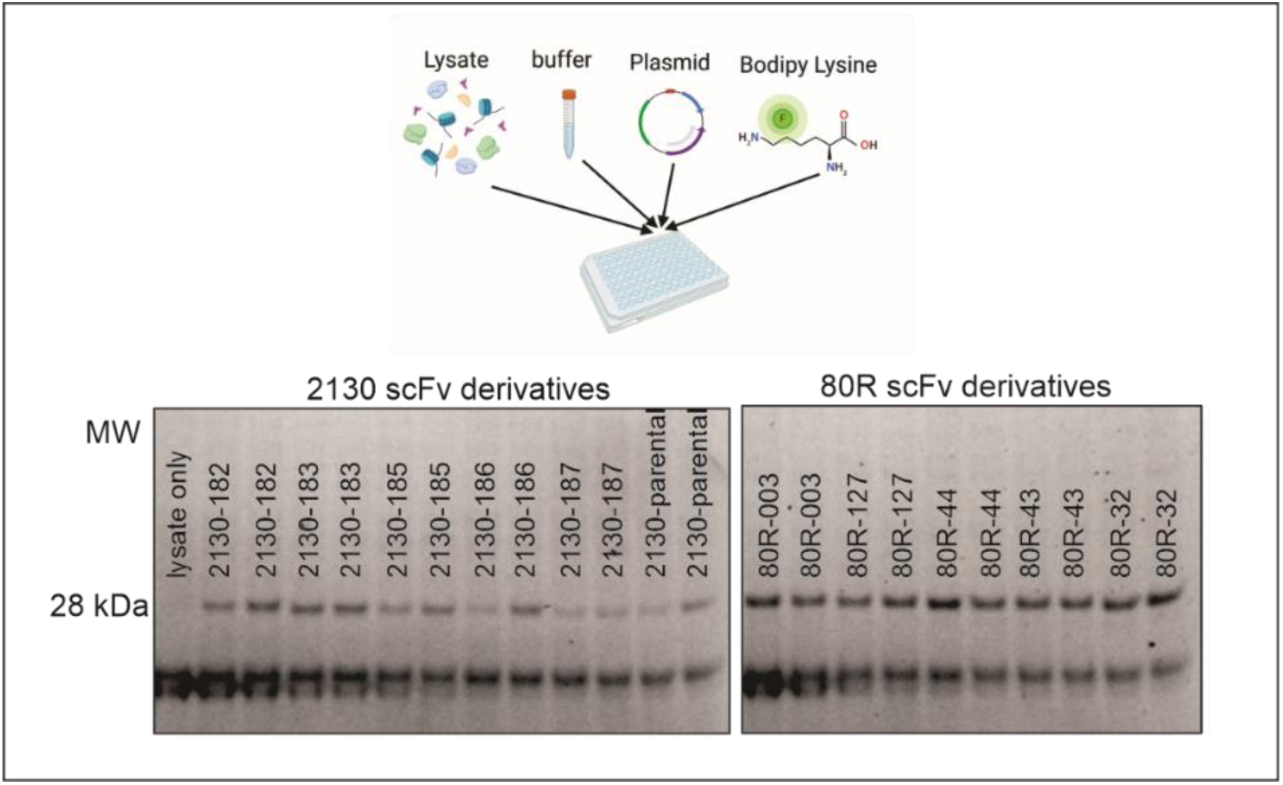
Expression of scFvs in *E. Coli*-Based Cell-Free Protein Synthesis (CFPS) and Visualization of Solubility Using Fluorescent Labeling. (Top): Schematic representation of the CFPS reaction setup, including lysate, reaction buffer, plasmid DNA, and BODIPY-Lysine-tRNA complex for in situ fluorescent labeling of newly synthesized proteins. **(Bottom**): SDS-PAGE gel showing CFPS-produced scFvs imaged at 488 nm without purification. The fluorescence signal confirms expression; however, analysis reveals low solubility across all tested variants, with 2130 scFvs exhibiting lower solubility than 80R derivatives. A consistent lower band around 18 kDa is observed across all reactions and is attributed to the BODIPY-lysine-tRNA complex. These results highlight the intrinsic solubility challenges of scFv expression in *E. coli* lysate-based CFPS and emphasize the need for chaperone-assisted folding (DnaK, DsbC) or post-expression refolding strategies to recover functional scFvs.

Given these limitations, achieving higher solubility required either the addition of chaperones (PUREfrex DnaK mix) and disulfide-bond isomerase enzymes (DsbC, PUREfrex) during CFPS (63–65), or the implementation of refolding techniques to recover functional scFvs from insoluble fractions (66). The limited solubility observed in the CFPS reactions underscores the inherent challenges of expressing scFvs in an *E. coli*-based cell-free system and highlights the necessity of solubility-enhancing strategies to obtain functional scFvs (60–62). These strategies include the addition of molecular chaperones (PUREfrex DnaK mix) and disulfide bond-forming enzymes (DsbC, PUREfrex), during translation, with concentrations suggested by the manufacturer (60–62). Alternatively, refolding techniques can be used to recover functional proteins from insoluble aggregates, which would require no addition of extra, potentially expensive, enzymes. Therefore, we to assess different strategies to improve scFv solubility yield.

### Optimized Cell-Free Production and Refolding Enhances scFv Binding Performance

To optimize scFv recovery and functionality, we selected two variants—2.59, the parental antibody of the 80R derivative lineage, and 80R-2.43, a strong binder identified in full-length IgG binding assays—for initial evaluation. Soluble yields from cell-free expression were initially low (∼0.02 mg/mL), with most of the product forming disulfide-linked dimers that did not exhibit target binding in functional assays (Figure 5 a, dimer band). To improve folding during expression, we supplemented reactions with DsbC, a disulfide isomerase, and the DnaK chaperone system, both of which modestly increased soluble yield up to 0.2 mg/mL (Figure 5a). However, the high cost of these folding additives limited their use for larger-scale production. To overcome this, we developed a two-stage refolding protocol aimed at enhancing both yield and activity of scFv proteins post-expression (Figure 5b). This refolding approach significantly improved recovery of functional protein, yielding up to 0.5 mg/mL of correctly folded scFv (Figure 5c). Importantly, the refolded products retained or improved their ability to bind WT RBD, as confirmed by SDS-PAGE and functional assays (Figure 6c), demonstrating that the two-stage refolding strategy is a cost-effective and scalable alternative to additive-based soluble expression.

**Figure 5.**
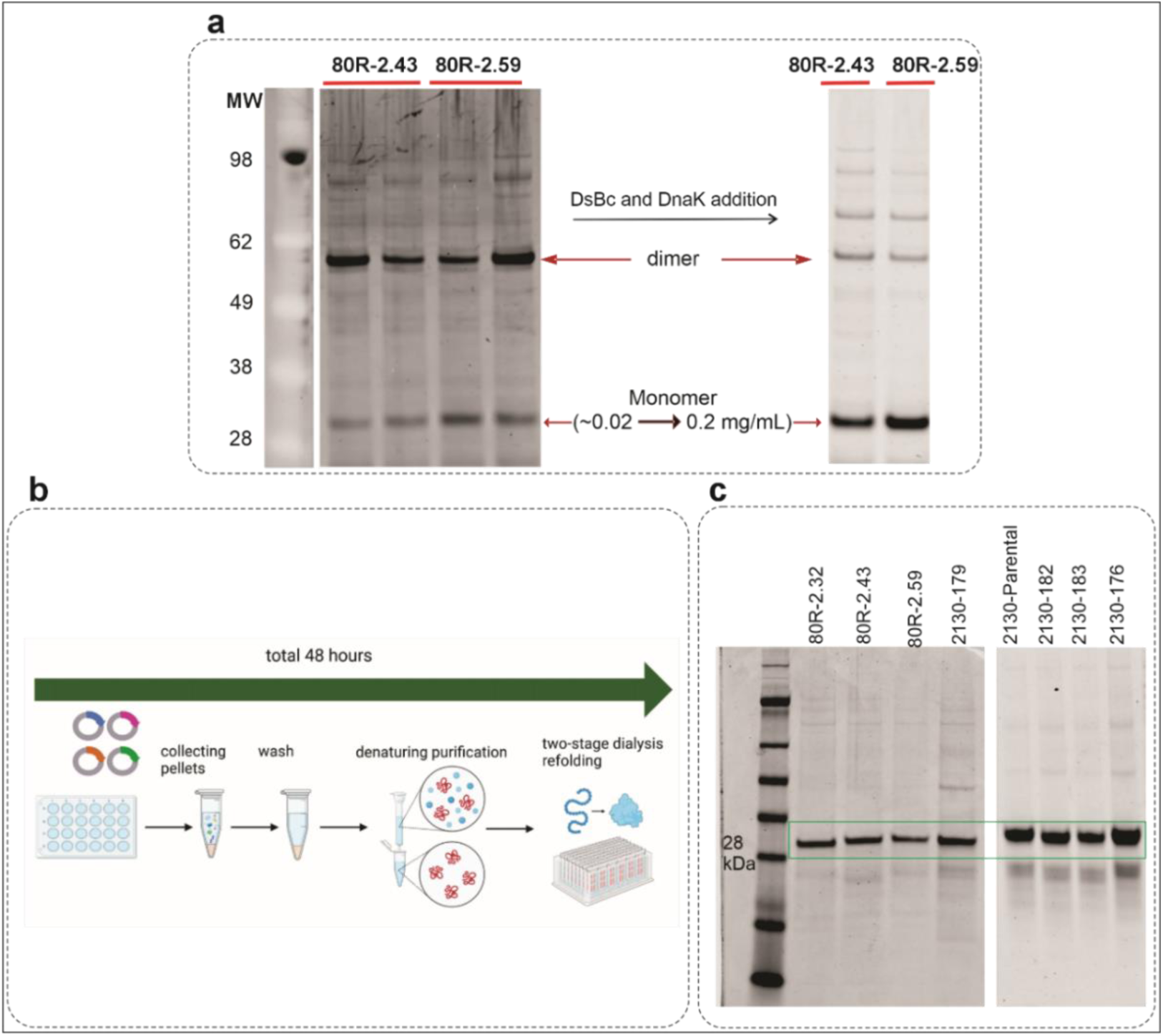
Optimization of scFv Production: Additives Improve Solubility in Cell-Free Expression, While Two-Stage Refolding Enables High-Yield Purification. **(a)** Two variants of scFvs, 80R-2.43 and 80R-2.59, were expressed in an E. coli-based cell-free protein synthesis (CFPS) system and subsequently purified, without additives, yielding minimal amounts of monomeric soluble protein. The addition of DsbC (disulfide isomerase) and DnaK (chaperone) to the reactions significantly increased the monomeric scFv yield, from 0.02 mg/mL to 0.2 mg/mL. **(b)** An illustration of the two-stage refolding process, developed to enhance cost-efficiency for scFvs expressed as inclusion bodies, is presented. This strategy substantially improved production yields. **(c)** The purification gel for eight scFv variants produced using the two-stage refolding workflow shows high monomeric yields of 0.5 mg/mL. The refolded scFvs retained structural integrity, solubility, and functionality, demonstrating the scalability of this approach for large-scale production.

**Figure 6.**
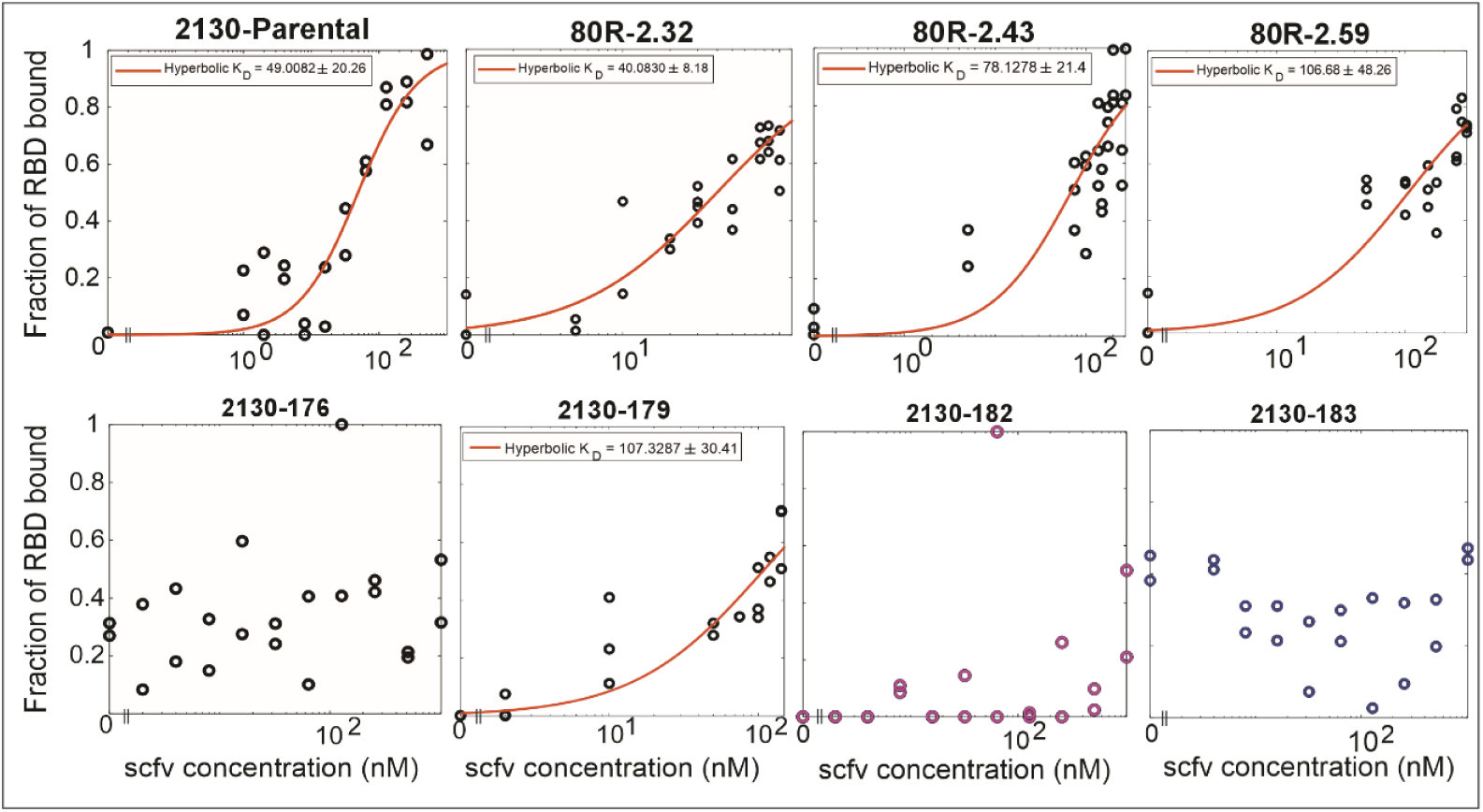
Binding Affinity Characterization of Refolded scFv Variants Reveals Specific Recognition of WT-RBD. Out of 8 tested scFvs, 5 were able to generate affinity binding data to WT-RBD. The fraction of RBD bound to scFv variants was determined by a two-species fit (equation 3) to the correlation curves. The hyperbolic binding equation (Equation 5) was fitted to the binding data to obtain the dissociation constant (K_D_s between 40-110 nM).

### FCS Enables Precise Measurement of scFv Binding Affinities

Following the successful validation of fluorescence correlation spectroscopy (FCS) for measuring antibody-antigen interactions, we applied FCS to quantify the binding affinities of anti-spike scFv variants produced using our in-house *E. coli* cell-free protein synthesis (CFPS) system. Among the eight scFv variants tested, we observed varying binding affinities. Three variants, 2130-parental, 80R-2.32, and 80R-2.43, demonstrated relatively strong binding, with respective *K*_D_s: 49.01 nM, 40.08 nM, and 78.12 nM, while 80R-3.59 and 2130-179 exhibited weaker affinities, with K_D_s equal to 106.06 nM and 107.32 nM. The *K*_D_ values for these 8 scFv variants and WT-RBD is listed in the table 2 and the binding curves are shown in figure 6. The remaining three variants did not show measurable binding within the nanomolar range. The binding affinities of 80R variant IgGs, produced in mammalian cell-based expression system, by ELISA technique is listed in the supplementary material. These findings highlight the variability in binding performance among the scFvs, emphasizing the need for optimization of production and folding processes to enhance functional yields.

**Table 2.** Dissociation Constants (*K*_*D*_) of scFv Variants for Wild-Type SARS-CoV-2 RBD.

| scFV | 2130-parental | 80R-2.32 | 80R-2.43 | 80R-2.59 | 2130-176 | 2130-179 | 2130-182 | 2130-183 |
| --- | --- | --- | --- | --- | --- | --- | --- | --- |
| $K_D$ (nM) | 49.01±20.26 | 40.08±8.18 | 78.1278±21.4 | 106.068±48.26 | - | 107.32±30.41 | - | - |

Binding affinities of scFvs prepared using both soluble expression and refolding methods were quantified using FCS. The scFvs produced via the refolding approach exhibited consistent binding profiles, with K_D_s comparable to those of scFvs generated in the soluble format and also in cell-based expression (supplementary material). Measurements were conducted using fluorescently labeled WT-RBD, labeled with Alexa647, as the constant antigen, with scFv concentrations titrated to generate binding curves. The Binding curves for the 8 tested scFv variants are demonstrated in figure 6. Shifts in FCS correlation curves reflected binding events (supplementary), with diffusion times increasing in the presence of scFvs. Hyperbolic fitting of the binding curves enabled precise determination of K_D_ values (table 2), confirming that the refolding process yields functional scFvs suitable for downstream applications. These results establish the two-stage refolding method as a reliable approach for producing high-quality scFvs.

### Efficient Cell-Free Production and Functional Validation of Fab Fragments

Given the limitations of full-length IgG production in bacterial CFPS systems, we next focused on the expression of antibody fragments called Fabs, that are structurally simpler than IgGs but retain antigen-binding functionality and are more similar to IgG structure than ScFvs. Fab fragments offer a compelling balance between structural integrity and reduced complexity, making them attractive candidates for rapid screening and functional evaluation. We therefore assessed the feasibility of producing soluble and functional Fab fragments using our *E. coli*-based CFPS platform. To test this, we utilized our *E. col*i-based CFPS platform to produce soluble and active 2130-Fab fragments with high efficiency. By co-expressing light and heavy chain DNA plasmids in a 1:1 molar ratio (30 nM total), we obtainedthe highest yields at approximately 0.5 mg/mL of purified Fab fragments (Figure 7a).

**Figure 7.**
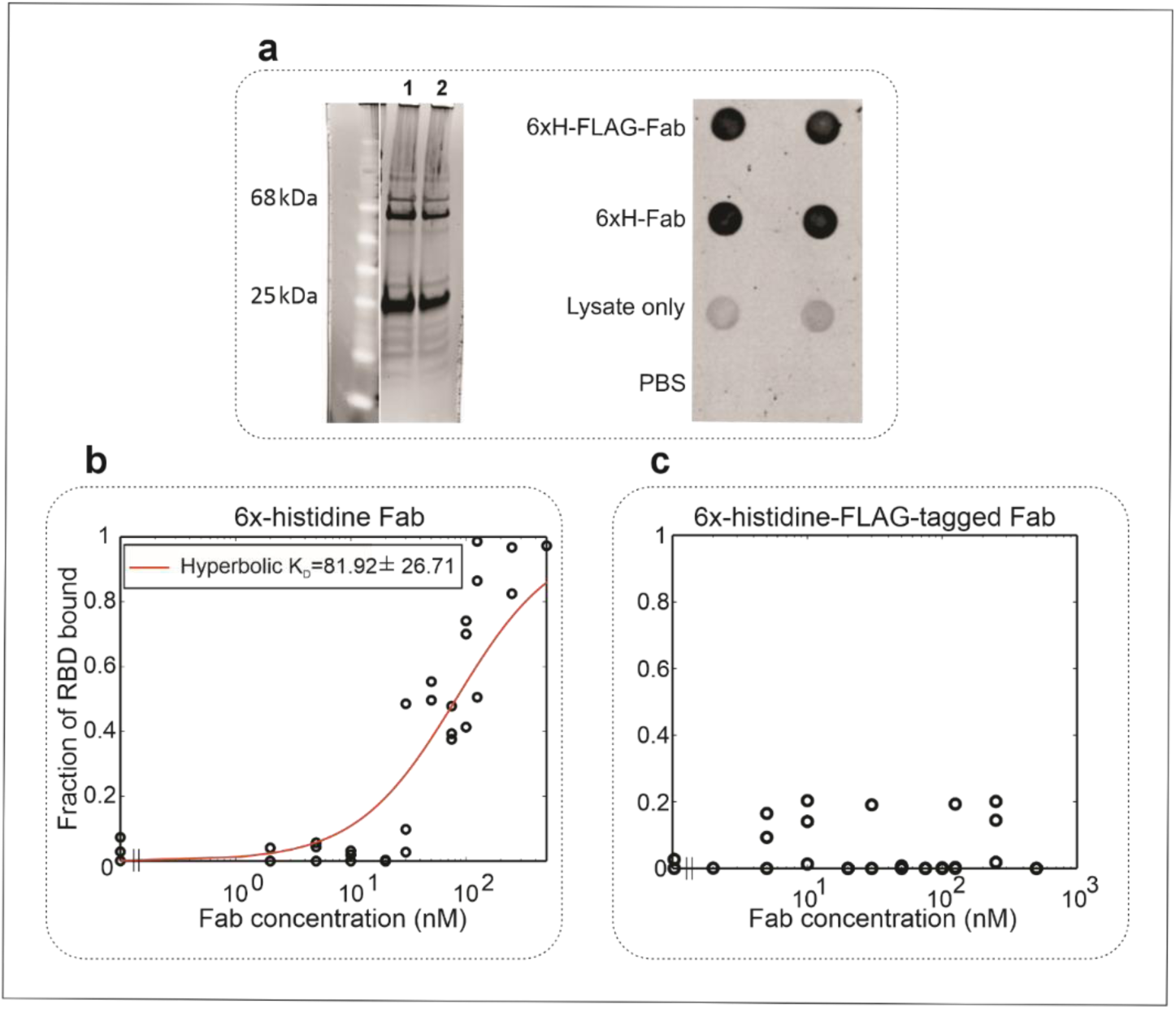
High-Yield Cell-free Production of Functional Fab Shows FLAG Tag Interferes with RBD Binding, while 6xHis Tag Does Not. **(a)** Reducing SDS-PAGE analysis of CFPS-produced 2130 Fab fragments purified via Ni-NTA. Lane 1 shows the 6x-histidine-tagged light chain and the heavy chain of 2130 Fab. Lane 2 contains the 6x-histidine-FLAG-tagged light chain along with the heavy chain of 2130 Fab. The lower band (∼25 kDa) represents dissociated heavy and light chains, while the upper band corresponds to the intact Fab fragment, composed of both chains. A dot blot assay (right) was used to confirm Fab structural integrity using a Fab-specific primary antibody which selectively recognizes conformational epitopes within the CH1 and CL domains. **(b)** Fraction of RBD bound to 6xHis-Fab was determined by a two species fit (eq. 6) to the correlation curves. The hyperbolic binding equation (Eq. 8) was fitted to the binding data to obtain the dissociation constant (*K*_*D*_ = 82 nM). **(c)** FLAG-tagged Fab construct exhibited no detectable RBD binding, suggesting steric hindrance or conformational interference.

To confirm the functionality of the CFPS-produced Fabs, we employed a dot blot assay using a conformational Fab-specific antibody (Sigma I5260), as this antibody selectively recognizes conformational epitopes within the CH1 and CL domains that are only exposed in correctly folded Fab structures, rather than linear or denatured epitopes. Detection of the Fabs by this antibody verified their proper folding and structural integrity, providing proof of concept to produce active Fabs using our CFPS platform (figure 7a). Two Fab constructs were tested: both shared identical heavy chains but differed in their light chain design, with one construct incorporating a FLAG tag alongside a His tag. While the FLAG tag did not affect solubility, yield, or its affinity to the Fab-specific antibody, its potential impact on its binding properties to RBD of the SARS-CoV-2 spike protein warranted further investigation. These results highlight the scalability and robustness of CFPS systems in generating antibody fragments suitable for multiple downstream applications.

### Cell-Free Produced Fab Fragments Exhibit High-Affinity Binding in FCS Assays

To assess binding affinities, fluorescence correlation spectroscopy (FCS) using the EI-FLEX instrument was performed using Alexa 647-labeled RBD (receptor-binding domain) of the SARS-CoV-2 spike protein. The reason for switching to Alexa647 dye was to match the red laser available on the EI-FLEX. Figure 7 (b and c) shows fraction of RBD bound to Fab when the concentration of labeled RBD was kept constant at around 0.5 nM and the concentration of Fab was increased. The 6xHis-tagged Fab construct demonstrated moderate binding to the RBD, with an equilibrium dissociation constant (K_D_) of approximately 80 nM (Figure 7b). However, the FLAG-tagged Fab construct, despite its high solubility and yield, exhibited no detectable binding to the RBD (Figure 7c). This finding suggests that the FLAG tag may interfere with structural conformation or epitope accessibility, emphasizing the importance of careful construct design in cell-free antibody production. Although the binding affinity of His-tagged Fab to WT-RBD was ∼10-fold lower than full-length IgG (0.72 nM compared to 80 nM), the ability for the Fab to bind to WT RBD in a measurable quantity suggests the easier-to-produce Fabs could be used to replace full-length IgGs in diagnostic settings. Furthermore, these results demonstrate the utility of CFPS for rapidly producing functional Fabs while underscoring the critical need for optimization to preserve binding activity.

## DISCUSSION

This study demonstrates significant advancements in antibody fragment production and characterization. One major strength lies in the successful application of fluorescence correlation spectroscopy (FCS) as a sensitive, high-throughput method for measuring antigen-antibody binding affinities. FCS was shown to be robust in detecting interactions across a range of binding strengths and antigen variants, enabling precise dissociation constant (*K*_D_) measurements at sub-nanomolar levels (67,68). Another key advantage is the use of cell-free protein synthesis (CFPS) platforms, which provide a rapid and efficient means for producing functional antibody fragments such as scFvs and Fabs (31,69). The study’s implementation of a two-stage refolding protocol further enhanced the solubility and yields of scFvs, making this approach scalable for larger production demands. Additionally, the CFPS system efficiently produced Fab fragments with proper folding and functional binding properties, highlighting its versatility for therapeutic and diagnostic applications to replace the more costly and time consuming IgGs. These innovations underscore the potential of integrating CFPS and FCS for accelerating antibody discovery and screening workflows as detailed below.

This study successfully validated FCS as a robust and reliable technique for quantifying antigen-antibody interactions. Through analysis of monoclonal antibodies wild-type2130 and S309, we demonstrated FCS’s ability to distinguish binding affinities to various RBD variants (56,57). The high sensitivity of FCS enabled the measurement of sub-nanomolar dissociation constants for wild-type2130 IgG binding to wild-type RBD. Conversely, the lack of detectable binding of wild-type2130 to BQ.1 and BQ.1.1 RBD variants underscores the impact of mutations on antibody recognition. S309 IgG exhibited broader binding capabilities, with measurable affinities to both wild-type and mutant RBD variants, highlighting its versatility. These results confirm FCS’s potential as a high-throughput platform for precise and sensitive kinetic analysis in antibody and antigen development workflows.

Binding affinities of refolded and soluble scFvs and Fabs were quantified also using FCS, which revealed consistent binding profiles between the two production methods. The refolded scFvs exhibited comparable K_D_s to those produced through soluble expression, confirming the reliability of the refolding process. The hyperbolic fitting of FCS-derived binding curves allowed precise determination of K_D_ values, establishing FCS as an indispensable tool for evaluating the functional quality of both scFvs and Fabs produced in cell-free systems. These findings demonstrate the compatibility of FCS with the future scale up of high-throughput antibody fragment screening. The variability in binding among the scFvs highlights the importance of incorporating folding processes in generating functional antibody fragments. Compared to soluble expression alone, the refolding approach increased overall recovery by 10-fold, as shown in analysis of results in figure 5. The ability of FCS to differentiate between high and low affinity provides a valuable tool for screening and optimizing scFv candidates for downstream applications. Applications include drug development, antibody design, and enzyme kinetics. They provide quantitative insights into binding properties, enabling advancements in therapeutic optimization, disease mechanism studies, and the development of diagnostic and research tools.

Despite its strengths, the study has certain limitations that warrant further attention. While FCS proved effective in measuring binding affinities, the technique requires access to specialized equipment and fluorescent labeling of antigens, which may limit its accessibility in resource-constrained settings. Additionally, variability in scFv binding affinities highlights the need for further optimization and specific protocols for production and folding of proteins, as three variants failed to exhibit measurable binding over the range of reagents studied. The presence of a FLAG tag in Fab constructs led to a complete loss of binding activity in one case, emphasizing the sensitivity that construct design may alter functional outcomes. This finding suggests that minor modifications, such as tags or fusion partners, could inadvertently interfere with functionality, necessitating thorough pre-screening of designs (70). Lastly, while CFPS systems demonstrated strong performance for scFv and Fab production, their applicability to full-length IgG synthesis or more complex antibody formats remains unexplored and may require more complex cell-free systems (71). Which highlights the need for more robust mammalian cell-free lysates. Addressing these limitations could further broaden the impact and utility of the CFPS methodologies described in this study.

### Future Directions and Applications

The results of this study highlight the utility of FCS as a versatile tool for measuring binding kinetics combined with the robustness of CFPS in generating different forms of human relevant functional antibody fragments. The two-stage refolding method offers a novel scalable solution for enhancing yields and maintaining activity of scFvs, while Fab production demonstrated the feasibility of rapidly generating antibody fragments with therapeutic and diagnostic potential. Future work will focus on further optimizing CFPS systems for a broader range of antibody formats and investigating structural and post translational factors affecting binding affinities (72,73). Together, these advancements position CFPS and FCS as critical tool for accelerating antibody discovery and development in both research and clinical contexts.

## CONCLUSIONS

This study demonstrates the efficacy of combining CFPS and FCS for the efficient production and characterization of antibody fragments. The *E. coli*-based CFPS system proved to be a reliable platform for generating scFvs and Fab fragments, with optimization strategies, such as the incorporation of folding enhancers and two-stage refolding protocols, significantly improving yields and solubility. The use of FCS enabled precise determination of binding kinetics, showcasing its potential as a high-throughput method for evaluating antibody-antigen interactions. These findings emphasize the scalability and flexibility of CFPS systems for therapeutic antibody development while highlighting FCS as a powerful tool for biophysical characterization. Future work will focus on expanding these methodologies to produce and evaluate more complex antibody formats, further solidifying this integrated approach as a cornerstone for rapid antibody discovery and screening in both research and clinical settings.

## ACKNOWLEGMENTS

The authors would like to thank Sean Gilmore and Brent Segelke for initial design and testing of the scFv plasmids; and Yaqing Wang, Sun Kyung Kim, and Bonnee Rubinfeld for providing the RBD and full-length antibody 98proteins. This project was funded by Lawrence Livermore National Laboratory Laboratory-Directed Research and Development grant 21ERD039 and partially supported by both the GUIDE and the ARPAH De Novo design of biologics (DeNOVO; AAH24039) programs. The GUIDE program is executed by the Joint Program Executive Office for Chemical, Biological, Radiological, and Nuclear Defense (JPEO-CBRND) Joint Project Lead for Enabling Biotechnologies (JPL CBRND EB) on behalf of the Department of Defense’s Chemical and Biological Defense Program. This effort was in collaboration with the Defense Health Agency (DHA) COVID funding initiative. Disclaimer: The views expressed in this paper reflect the views of the authors and do not necessarily reflect the position of the Department of the Army, Department of Defense, nor the United States Government. References to non-federal entitiesdo not constitute nor imply Department of Defense or Army endorsement of any company or organization. This work performed under the auspices of the U.S. Department of Energy by Lawrence Livermore National Laboratory under Contract DE-AC52-07NA27344.

## REFERENCES

1. Kim TK, Eberwine JH. Mammalian cell transfection: the present and the future. Anal Bioanal Chem. 2010 Aug 1;397(8):3173–8.

2. Geisse S, Henke M. Large-scale Transient Transfection of Mammalian Cells: A Newly Emerging Attractive Option for Recombinant Protein Production. J Struct Funct Genomics. 2005 Sep 1;6(2):165–70.

3. Rosser MP, Xia W, Hartsell S, McCaman M, Zhu Y, Wang S, et al. Transient transfection of CHO-K1-S using serum-free medium in suspension: a rapid mammalian protein expression system. Protein Expr Purif. 2005 Apr 1;40(2):237–43.

4. Katzen F, Chang G, Kudlicki W. The past, present and future of cell-free protein synthesis. Trends Biotechnol. 2005 Mar 1;23(3):150–6.

5. Carlson ED, Gan R, Hodgman CE, Jewett MC. Cell-free protein synthesis: Applications come of age. Biotechnol Adv. 2012 Sep;30(5):1185–94.

6. Zemella A, Thoring L, Hoffmeister C, Kubick S. Cell-Free Protein Synthesis: Pros and Cons of Prokaryotic and Eukaryotic Systems. ChemBioChem. 2015;16(17):2420–31.

7. Jin X, Hong SH. Cell-free protein synthesis for producing ‘difficult-to-express’ proteins. Biochem Eng J. 2018 Oct 15;138:156–64.

8. Sawasaki T, Ogasawara T, Morishita R, Endo Y. A cell-free protein synthesis system for high-throughput proteomics. Proc Natl Acad Sci. 2002 Nov 12;99(23):14652–7.

9. Dondapati SK, Stech M, Zemella A, Kubick S. Cell-Free Protein Synthesis: A Promising Option for Future Drug Development. BioDrugs. 2020 Jun;34(3):327–48.

10. Nirenberg MW, Matthaei JH. The dependence of cell-free protein synthesis in E. coli upon naturally occurring or synthetic polyribonucleotides. Proc Natl Acad Sci. 1961 Oct;47(10):1588–602.

11. Weisser NE, Hall JC. Applications of single-chain variable fragment antibodies in therapeutics and diagnostics. Biotechnol Adv. 2009 Jul 1;27(4):502–20.

12. Monnier PP, Vigouroux RJ, Tassew NG. In Vivo Applications of Single Chain Fv (Variable Domain) (scFv) Fragments. Antibodies. 2013 Jun;2(2):193–208.

13. Gezehagn Kussia G, Tessema TS. The Potential of Single-Chain Variable Fragment Antibody: Role in Future Therapeutic and Diagnostic Biologics. J Immunol Res. 2024;2024(1):1804038.

14. Crivianu-Gaita V, Thompson M. Aptamers, antibody scFv, and antibody Fab’ fragments: An overview and comparison of three of the most versatile biosensor biorecognition elements. Biosens Bioelectron. 2016 Nov 15;85:32–45.

15. Bever CS, Dong JX, Vasylieva N, Barnych B, Cui Y, Xu ZL, et al. VHH antibodies: emerging reagents for the analysis of environmental chemicals. Anal Bioanal Chem. 2016 Sep;408(22):5985–6002.

16. Stech M, Hust M, Schulze C, Dübel S, Kubick S. Cell-free eukaryotic systems for the production, engineering, and modification of scFv antibody fragments. Eng Life Sci. 2014 Jul;14(4):387–98.

17. Flanagan RJ, Jones AL. Fab Antibody Fragments: Some Applications in Clinical Toxicology. Drug Saf. 2004;27(14):1115–33.

18. Stafford RL, Matsumoto ML, Yin G, Cai Q, Fung JJ, Stephenson H, et al. In vitro Fab display: a cell-free system for IgG discovery. Protein Eng Des Sel. 2014 Apr 1;27(4):97– 109.

19. Malpiedi LP, Díaz CA, Nerli BB, Pessoa A. Single-chain antibody fragments: Purification methodologies. Process Biochem. 2013 Aug 1;48(8):1242–51.

20. Holliger P, Hudson PJ. Engineered antibody fragments and the rise of single domains. Nat Biotechnol. 2005 Sep;23(9):1126–36.

21. Bates A, Power CA. David vs. Goliath: The Structure, Function, and Clinical Prospects of Antibody Fragments. Antibodies. 2019 Jun;8(2):28.

22. Flanagan RJ, Jones AL. Fab Antibody Fragments. Drug Saf. 2004 Dec 1;27(14):1115–33.

23. Zemella A, Thoring L, Hoffmeister C, Kubick S. Cell-Free Protein Synthesis: Pros and Cons of Prokaryotic and Eukaryotic Systems. ChemBioChem. 2015;16(17):2420–31.

24. Stech M, Nikolaeva O, Thoring L, Stöcklein WFM, Wüstenhagen DA, Hust M, et al. Cell-free synthesis of functional antibodies using a coupled in vitro transcription-translation system based on CHO cell lysates. Sci Rep. 2017 Sep 20;7(1):12030.

25. Kanter G, Yang J, Voloshin A, Levy S, Swartz JR, Levy R. Cell-free production of scFv fusion proteins: an efficient approach for personalized lymphoma vaccines. Blood. 2007 Apr 15;109(8):3393–9.

26. Yin G, Garces, Eudean D., Yang, Junhao, Zhang, Juan, Tran, Cuong, Steiner, Alexander R, et al. Aglycosylated antibodies and antibody fragments produced in a scalable in vitro transcription-translation system. mAbs. 2012 Mar 1;4(2):217–25.

27. Goerke AR, Swartz JR. High-level cell-free synthesis yields of proteins containing site-specific non-natural amino acids. Biotechnol Bioeng. 2009;102(2):400–16.

28. Ban B, Sharma M, Shetty J. Optimization of Methods for the Production and Refolding of Biologically Active Disulfide Bond-Rich Antibody Fragments in Microbial Hosts. Antibodies. 2020 Aug 5;9(3):39.

29. Butler JE. Enzyme-Linked Immunosorbent Assay. J Immunoassay [Internet]. 2000 May 1 [cited 2025 Feb 9]; Available from: https://www.tandfonline.com/doi/abs/10.1080/01971520009349533

30. Fischer SK, Joyce A, Spengler M, Yang TY, Zhuang Y, Fjording MS, et al. Emerging Technologies to Increase Ligand Binding Assay Sensitivity. AAPS J. 2015 Jan 1;17(1):93– 101.

31. Hunt AC, Vögeli B, Hassan AO, Guerrero L, Kightlinger W, Yoesep DJ, et al. A rapid cell-free expression and screening platform for antibody discovery. Nat Commun. 2023 Jul 3;14(1):3897.

32. Designing binding kinetic assay on the bio-layer interferometry (BLI) biosensor to characterize antibody-antigen interactions-ScienceDirect [Internet]. [cited 2025 Apr 8]. Available from: https://www.sciencedirect.com/science/article/pii/S0003269717303342?via%3Dihub

33. Ruan Q, Tetin SY. Applications of dual-color fluorescence cross-correlation spectroscopy in antibody binding studies. Anal Biochem. 2008 Mar 1;374(1):182–95.

34. Kuroki K, Kobayashi S, Shiroishi M, Kajikawa M, Okamoto N, Kohda D, et al. Detection of weak ligand interactions of leukocyte Ig-like receptor B1 by fluorescence correlation spectroscopy. J Immunol Methods. 2007 Mar 30;320(1):172–6.

35. Tetin SY, Ruan Q, Saldana SC, Pope MR, Chen Y, Wu H, et al. Interactions of Two Monoclonal Antibodies with BNP: High Resolution Epitope Mapping Using Fluorescence Correlation Spectroscopy. Biochemistry. 2006 Nov 1;45(47):14155–65.

36. Wu CY, Huang CK, Chung CY, Huang IP, Hwu Y, Yang CS, et al. Probing the binding kinetics of proinflammatory cytokine–antibody interactions using dual color fluorescence cross correlation spectroscopy. Analyst. 2011 Apr 25;136(10):2111–8.

37. Barbero N, Napione L, Quagliotto P, Pavan S, Barolo C, Barni E, et al. Fluorescence anisotropy analysis of protein–antibody interaction. Dyes Pigments. 2009 Nov 1;83(2):225– 9.

38. Liu C, Hoang-Phou SA, Ye C, Laurence EJ, Laurence MJ, Fong EJ, et al. Real-Time Affinity Measurements of Proteins Synthesized in Cell-Free Lysate Using Fluorescence Correlation Spectroscopy. Anal Chem. 2025 May 13;97(18):9638–47.

39. Magde D, Elson E, Webb WW. Thermodynamic Fluctuations in a Reacting System— Measurement by Fluorescence Correlation Spectroscopy. Phys Rev Lett. 1972 Sep 11;29(11):705–8.

40. Rigler R, Mets □., Widengren J, Kask P. Fluorescence correlation spectroscopy with high count rate and low background: analysis of translational diffusion. Eur Biophys J [Internet]. 1993 Aug [cited 2025 Jan 19];22(3). Available from: http://link.springer.com/10.1007/BF00185777

41. Sharma A, Sarkar A, Goswami D, Bhattacharyya A, Enderlein J, Kumbhakar M. Determining Metal Ion Complexation Kinetics with Fluorescent Ligands by Using Fluorescence Correlation Spectroscopy. ChemPhysChem. 2019;20(16):2093–102.

42. Porciani D, Alampi MM, Abbruzzetti S, Viappiani C, Delcanale P. Fluorescence Correlation Spectroscopy as a Versatile Method to Define Aptamer–Protein Interactions with Single-Molecule Sensitivity. Anal Chem. 2024 Jan 9;96(1):137–44.

43. Beam M, Silva MC, Morimoto RI. Dynamic Imaging by Fluorescence Correlation Spectroscopy Identifies Diverse Populations of Polyglutamine Oligomers Formed in Vivo. J Biol Chem. 2012 Jul 27;287(31):26136–45.

44. Yu L, Lei Y, Ma Y, Liu M, Zheng J, Dan D, et al. A Comprehensive Review of Fluorescence Correlation Spectroscopy. Front Phys [Internet]. 2021 Apr 12 [cited 2025 Apr 8];9. Available from: https://www.frontiersin.org/journals/physics/articles/10.3389/fphy.2021.644450/full

45. Matsunaga R, Ujiie K, Inagaki M, Fernández Pérez J, Yasuda Y, Mimasu S, et al. High-throughput analysis system of interaction kinetics for data-driven antibody design. Sci Rep. 2023 Nov 21;13(1):19417.

46. Wang X, Phan MM, Sun Y, Koerber JT, Ho H, Chen Y, et al. Development of an SPR-based binding assay for characterization of anti-CD20 antibodies to CD20 expressed on extracellular vesicles. Anal Biochem. 2022 Jun 1;646:114635.

47. Liu C, Hoang-Phou SA, Ye C, Laurence EJ, Laurence MJ, Fong EJ, et al. Real-time affinity measurements of proteins synthesized in cell-free lysate using fluorescence correlation spectroscopy [Internet]. bioRxiv; 2024 [cited 2025 Jan 17]. p. 2024.09.28.615564. Available from: https://www.biorxiv.org/content/10.1101/2024.09.28.615564v1

48. Desautels TA, Arrildt KT, Zemla AT, Lau EY, Zhu F, Ricci D, et al. Computationally restoring the potency of a clinical antibody against Omicron. Nature. 2024 May;629(8013):878–85.

49. Principles of Fluorescence Spectroscopy | SpringerLink [Internet]. [cited 2025 Jan 22]. Available from: https://link.springer.com/book/10.1007/978-0-387-46312-4

50. Frontiers | A Comprehensive Review of Fluorescence Correlation Spectroscopy [Internet]. [cited 2025 Jan 22]. Available from: https://www.frontiersin.org/journals/physics/articles/10.3389/fphy.2021.644450/full

51. Banachowicz E, Patkowski A, Meier G, Klamecka K, Gapiński J. Successful FCS Experiment in Nonstandard Conditions. Langmuir. 2014 Jul 29;30(29):8945–55.

52. Diffusion of single molecules through a Gaussian laser beam-Astrophysics Data System [Internet]. [cited 2025 Jan 22]. Available from: https://ui.adsabs.harvard.edu/abs/1993SPIE.1921..239R/abstract

53. Pinto D, Park YJ, Beltramello M, Walls AC, Tortorici MA, Bianchi S, et al. Cross-neutralization of SARS-CoV-2 by a human monoclonal SARS-CoV antibody. Nature. 2020 Jul;583(7815):290–5.

54. Qu P, Evans JP, Faraone JN, Zheng YM, Carlin C, Anghelina M, et al. Enhanced neutralization resistance of SARS-CoV-2 Omicron subvariants BQ.1, BQ.1.1, BA.4.6, BF.7, and BA.2.75.2. Cell Host Microbe. 2023 Jan 11;31(1):9–17.e3.

55. Chen Y, Zhao X, Zhou H, Zhu H, Jiang S, Wang P. Broadly neutralizing antibodies to SARS-CoV-2 and other human coronaviruses. Nat Rev Immunol. 2023 Mar;23(3):189–99.

56. He Q, Wu L, Xu Z, Wang X, Xie Y, Chai Y, et al. An updated atlas of antibody evasion by SARS-CoV-2 Omicron sub-variants including BQ.1.1 and XBB. Cell Rep Med. 2023 Apr 18;4(4):100991.

57. de Campos-Mata L, Trinité B, Modrego A, Tejedor Vaquero S, Pradenas E, Pons-Grífols A, et al. A monoclonal antibody targeting a large surface of the receptor binding motif shows pan-neutralizing SARS-CoV-2 activity. Nat Commun. 2024 Feb 5;15(1):1051.

58. Pinto D, Park YJ, Beltramello M, Walls AC, Tortorici MA, Bianchi S, et al. Cross-neutralization of SARS-CoV-2 by a human monoclonal SARS-CoV antibody. Nature. 2020 Jul;583(7815):290–5.

59. Kohyama S, Frohn BP, Babl L, Schwille P. Machine learning-aided design and screening of an emergent protein function in synthetic cells. Nat Commun. 2024 Mar 5;15(1):2010.

60. Sandomenico A, Sivaccumar JP, Ruvo M. Evolution of Escherichia coli Expression System in Producing Antibody Recombinant Fragments. Int J Mol Sci. 2020 Jan;21(17):6324.

61. Zemella A, Thoring L, Hoffmeister C, Kubick S. Cell-Free Protein Synthesis: Pros and Cons of Prokaryotic and Eukaryotic Systems. Chembiochem. 2015 Nov;16(17):2420–31.

62. Liu M, Wang B, Wang F, Yang Z, Gao D, Zhang C, et al. Soluble expression of single-chain variable fragment (scFv) in Escherichia coli using superfolder green fluorescent protein as fusion partner. Appl Microbiol Biotechnol. 2019 Aug;103(15):6071–9.

63. Zhang Z, Li ZH, Wang F, Fang M, Yin CC, Zhou ZY, et al. Overexpression of DsbC and DsbG markedly improves soluble and functional expression of single-chain Fv antibodies in Escherichia coli. Protein Expr Purif. 2002 Nov;26(2):218–28.

64. Estabragh AM, Sadeghi HMM, Akbari V. Co-Expression of Chaperones for Improvement of Soluble Expression and Purification of An Anti-HER2 scFv in Escherichia Coli. Adv Biomed Res. 2022 Dec 26;11:117.

65. Wang R, Xiang S, Feng Y, Srinivas S, Zhang Y, Lin M, et al. Engineering production of functional scFv antibody in E. coli by co-expressing the molecule chaperone Skp. Front Cell Infect Microbiol [Internet]. 2013 Nov 6 [cited 2025 Apr 8];3. Available from: https://www.frontiersin.org/journals/cellular-and-infection-microbiology/articles/10.3389/fcimb.2013.00072/full

66. Geisbrecht BV, Bouyain S, Pop M. An optimized system for expression and purification of secreted bacterial proteins. Protein Expr Purif. 2006 Mar 1;46(1):23–32.

67. Porciani D, Alampi MM, Abbruzzetti S, Viappiani C, Delcanale P. Fluorescence Correlation Spectroscopy as a Versatile Method to Define Aptamer–Protein Interactions with Single-Molecule Sensitivity. Anal Chem. 2023 Dec 21;96(1):137–44.

68. Yu L, Lei Y, Ma Y, Liu M, Zheng J, Dan D, et al. A Comprehensive Review of Fluorescence Correlation Spectroscopy. Front Phys [Internet]. 2021 Apr 12 [cited 2025 Apr 9];9. Available from: https://www.frontiersin.org/journals/physics/articles/10.3389/fphy.2021.644450/full

69. Haueis L, Stech M, Kubick S. A Cell-free Expression Pipeline for the Generation and Functional Characterization of Nanobodies. Front Bioeng Biotechnol [Internet]. 2022 Apr 28 [cited 2025 Apr 10];10. Available from: https://www.frontiersin.org/journals/bioengineering-and-biotechnology/articles/10.3389/fbioe.2022.896763/full

70. Krachmarova E, Tileva M, Lilkova E, Petkov P, Maskos K, Ilieva N, et al. His-FLAG Tag as a Fusion Partner of Glycosylated Human Interferon-Gamma and Its Mutant: Gain or Loss? BioMed Res Int. 2017;2017:3018608.

71. Krishna S, Jung, Sang Taek, and Lee EY. Escherichia coli and Pichia pastoris: microbial cell-factory platform for-full-length IgG production. Crit Rev Biotechnol. 2025 Jan 2;45(1):191–213.

72. Williams AJ, Warfel KF, Desai P, Li J, Lee JJ, Wong DA, et al. A low-cost recombinant glycoconjugate vaccine confers immunogenicity and protection against enterotoxigenic Escherichia coli infections in mice. Front Mol Biosci. 2023;10:1085887.

73. Kightlinger W, Duncker KE, Ramesh A, Thames AH, Natarajan A, Stark JC, et al. A cell-free biosynthesis platform for modular construction of protein glycosylation pathways. Nat Commun. 2019 Nov 27;10(1):5404.

